# Sequence charge decoration organizes salt response regimes in intrinsically disordered proteins: an interpretable machine-learning study

**DOI:** 10.64898/2026.06.01.729380

**Authors:** Mahesh Aryal

## Abstract

The conformational ensembles of intrinsically disordered proteins (IDPs) respond sensitively to ionic strength, with the direction and magnitude of response varying widely across sequence classes from polyelectrolyte contraction to polyampholyte swelling. Recent sequence-conditioned models provide rapid access to full ensembles or ensemble-averaged properties at specified solution conditions, but do not directly identify which low-dimensional polymer physics descriptors organize salt response behavior. Here, we construct a 511-sequence library spanning controlled *κ*-variants, NCPR series, IDRome-stratified natural IDRs, and low-FCR IDRome sequences, and perform 2,555 CALVADOS-2 simulations across five monovalent salt concentrations (50 mM to 500 mM). For each sequence, we extract the salt-response slope, d*R*_g_*/*d[salt], and assign one of four regimes: polyelectrolyte contraction, polyampholyte swelling, non-monotonic response, or salt-insensitive behavior. Using eight theory-motivated sequence descriptors, we find that sequence charge decoration weighted by chain length, SCD × *N*, is the dominant coordinate organizing salt response, accounting for ~40% of total SHAP attribution and exceeding the next feature by more than twice. Ridge regression explains substantial in-distribution variance (*R*^2^ = 0.83 under random cross-validation), whereas gradient-boosted trees improve in-distribution performance (*R*^2^ = 0.97) and retain predictive power under the more stringent leave-one-subset-out validation test (*R*^2^ = 0.60), indicating that salt response contains transferable but nonlinear sequence-encoded structure. Regime classification robustly recovers the direction of salt response, with no polyelectrolyte–polyampholyte confusion, whereas non-monotonic and salt-insensitive sequences remain harder to distinguish from static sequence features alone. Together, these results establish SCD × *N* as a compact, interpretable organizing coordinate for CALVADOS-2-derived IDP salt response and provide a polymer-physics, feature-level complement to ensemble-level generative models.

## 1 Introduction

Biopolymers such as proteins respond to their solvent environment by changing their conformations and overall dimensions as surrounding conditions vary. Their equilibrium dimensions reflect a balance between chain configurational entropy and effective segment–segment and segment–solvent interactions. Environmental variables that alter this balance can therefore reshape the chain through mechanisms such as excluded-volume and depletion interactions induced by cosolutes or electrostatic screening in charged polymers Asakura and Oosawa (1954); de Gennes (1979); Rubinstein and Colby (2003); Dobrynin et al. (1995); Aryal and Denton (2026). This environmental sensitivity is particularly pronounced in intrinsically disordered proteins (IDPs), which sample broad conformational ensembles rather than adopting a single folded structure Wright and Dyson (2015); van der Lee et al. (2014). Because many IDP sequences are highly charged or polyampholytic, intrachain electrostatic interactions strongly influence ensemble-averaged observables such as the radius of gyration *R*_g_, end-to-end distance *R*_ee_, and hydrodynamic radius *R*_*h*_. Changes in ionic strength screen these interactions and alter the effective excluded volume of the chain, producing measurable expansion, contraction, non-monotonic responses, or weak salt dependence Müller-Späth et al. (2010); Waszkiewicz et al. (2024); Cubuk and Soranno (2022); Banks et al. (2018). These conformational responses are not merely structural details: they can modulate phase separation, molecular recognition, and the stability and material properties of biomolecular condensates Lin et al. (2016); Wicky et al. (2017); Mittag and Pappu (2022). Moreover, cells are not maintained at a single ionic strength. IDPs encounter ionic variations across cellular compartments, during osmotic stress, and under the diverse solution conditions used in reconstituted experiments. Thus, how an IDP ensemble *responds* to changes in salt concentration—not merely its dimensions at a single condition—is itself a functionally relevant sequence-dependent property.

The theoretical foundation for understanding charge driven IDP conformations rests on classical polyelectrolyte and polyampholyte theory. Higgs and Joanny showed that randomly charged chains can exhibit salt-dependent swelling or collapse depending on the balance between net charge, charge fluctuations, and screening Higgs and Joanny (1991). Later, sequence specific theories connected these ideas more directly to IDPs: Das and Pappu introduced the charge patterning parameter *κ* to quantify the segregation of oppositely charged residues along a sequence, whereas Sawle and Ghosh introduced the sequence charge decoration (SCD) descriptor to capture the patterning-weighted electrostatic contribution to chain dimensions Das and Pappu (2013); Sawle and Ghosh (2015). While SCD is now well established as a predictor of IDP dimensions at fixed conditions, its quantitative role in governing *salt response* specifically, across broad sequence space, has not been systematically established. Random-phase-approximation approaches further showed that charge patterning can govern both single chain dimensions and the phase behavior of polyampholytic IDPs Lin et al. (2016). Extending these ideas to finite salt, Huihui, Firman, and Ghosh predicted that polyelectrolytic and polyampholytic sequences need not respond monotonically or even in the same direction to added salt, implying a sequence dependent taxonomy of response regimes Huihui et al. (2018). Coarse-grained molecular dynamics surveys by Wohl, Jakubowski, and Zheng subsequently emphasized that salt response can also include residue specific salting out contributions beyond simple Debye–Hückel screening Wohl et al. (2021). Together, these studies establish IDP salt response as a sequence-dependent polymer physics problem and predict that distinct response regimes should exist; what remains is to map that regime structure empirically across controlled sequence space. Large scale simulations and machine learning models have begun to make IDP ensemble characterization more accessible. CALVADOS-based simulations have enabled proteome scale prediction of IDP dimensions, including the IDRome database of conformational ensembles for tens of thousands of human IDRs, primarily at fixed physiological ionic strength Tesei et al. (2024). In parallel, sequence-to-property models such as ALBATROSS demonstrated that ensemble-averaged IDP observables can be predicted rapidly from amino acid sequence Lotthammer et al. (2024). More recently, STARLING introduced a latent diffusion generative framework for producing full coarse-grained IDR conformational ensembles from sequence, with conditioning on ionic strength and the ability to interpolate between salt conditions Novak et al. (2026). These advances show that sequence conditioned learning can capture important aspects of IDP ensemble physics. What remains missing is a reduced, interpretable account: which polymer physics descriptors govern the *direction* and *magnitude* of salt response, and how sequences organize into distinct response regimes across sequence space.

Here we take a complementary, response focused approach. Rather than generating full ensembles at individual salt concentrations, we construct a systematic 511-sequence CALVADOS-2 simulation library spanning five monovalent salt concentrations from 50 mM to 500 mM. Using a single coarse-grained model allows us to isolate sequence encoded electrostatic response within a controlled and internally consistent simulation framework. For each sequence, we compress the salt dependence of chain dimensions into a response slope, d*R*_g_*/*d[salt], and assign one of four polymer-physics regimes: polyelectrolyte contraction, polyampholyte swelling, non-monotonic response, or salt-insensitive behavior. Using ridge regression, gradient-boosted trees, and SHAP analysis on a compact set of theory motivated sequence descriptors, we show that SCD × *N* acts as the dominant coordinate organizing salt response. We further find that the *direction* of response is robustly encoded by these descriptors, whereas generalization across sequence families is the principal difficulty: linear models extrapolate poorly between families, and nonlinear feature interactions are required to recover predictive accuracy out of distribution. This work therefore provides an interpretable machine learning framework connecting sequence charge patterning to salt-response regimes, complementing recent sequence-to-ensemble models with a reduced polymer-physics map of electrostatic response.

## 2 Methods

### 2.1 Sequence library

We constructed a 511-sequence library designed to span a broad range of charge compositions, charge patterning architectures, and expected polymer physics salt response regimes. The library comprises four subsets: two synthetic subsets generated to provide controlled coverage of charge sequence space, and two natural subsets drawn from the IDRome database Tesei et al. (2024) to provide biological realism. Synthetic sequences were built on a neutral Gly/Ser/Ala/Pro background using lysine (K) and glutamate (E) as the positively and negatively charged residues, following the Das-Pappu construction Das and Pappu (2013).

#### *κ*-variants (195 sequences)

To isolate the effect of charge patterning at fixed composition, we generated polyampholyte sequences with near-zero net charge and systematically varied the charge-patterning parameter *κ* Das and Pappu (2013). For each combination of length *N* ∈ {50, 100, 200} and fraction of charged residues FCR ∈ {0.30, 0.50}, we placed equal numbers of K and E residues and accepted sequences into eleven target *κ* bins spanning 0.05–0.55 (three sequences per bin, tolerance |Δ*κ*| ≤ 0.04). Well-mixed configurations were obtained by random charge placement; highly segregated (high-*κ*) configurations were obtained by biased clustered placement to fill bins that random placement reaches only rarely. *κ* was evaluated with localCIDER Holehouse et al. (2017).

#### NCPR series (108 sequences)

To probe the transition from weakly charged chains to strong polyelectrolytes, we fixed FCR = 0.40 and titrated the net charge per residue NCPR across nine values from iyu −0.40 to +0.40 in steps of 0.10, at lengths *N* ∈ {50, 100, 200} (four sequences per cell). At the extremes (|NCPR| = 0.40, i.e. FCR = |NCPR|) all charged residues share the same sign, yielding pure polyelectrolytes; *κ* is undefined for these single-charge-type sequences and is set to zero in the feature matrix (Section 2.6).

#### IDRome stratified (150 sequences)

To provide biologically representative natural sequences, we sampled the IDRome database Tesei et al. (2024) (28,058 human IDRs with CALVADOS-2 properties precomputed at 150 mM). We restricted to sequences of length 30–300 residues containing only the twenty standard amino acids, then drew a sample stratified over a 5 × 5 grid of quantile bins in (SCD, *N*) (six sequences per cell), ensuring coverage of the corners of natural charge decoration and length space rather than only the population mode.

#### IDRome low-FCR (58 sequences)

To sample weakly charged, nearly salt-insensitive sequences, we filtered IDRome to FCR *<* 0.10, removed sequences already selected for the stratified subset, and sampled the remainder stratified over aromatic fraction and length. Truly low-charge IDRs are biologically uncommon (only 81 of the length-filtered IDRome entries satisfy FCR *<* 0.10), so this subset is intentionally smaller than the others.

For each sequence we computed a common set of composition and charge-patterning descriptors: sequence length *N*, fraction of charged residues FCR, net charge per residue NCPR, charge-patterning parameter *κ* Das and Pappu (2013), sequence charge decoration SCD Sawle and Ghosh (2015), mean hydropathy, and aromatic fraction. SCD quantifies the patterning-weighted electrostatic contribution to chain dimensions and is defined as Sawle and Ghosh (2015)

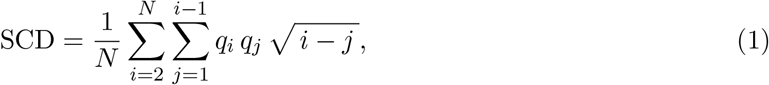

where *q*_*k*_ = +1 for K/R, −1 for D/E, and 0 otherwise, and the indices run over residue positions. For the synthetic subsets, FCR, NCPR, and *κ* were computed with localCIDER Holehouse et al. (2017) and SCD from Eq. (1); for the IDRome subsets we used the values precomputed in the database. Mean hydropathy was computed uniformly across all 511 sequences using the Kyte– Doolittle scale Kyte and Doolittle (1982) so that this feature is comparable between synthetic and natural subsets. Aromatic fraction and the CALVADOS hydropathy parameter *λ* were used to characterize the library but were not included in the final eight-feature model unless stated otherwise. These quantities were used both to characterize the library and as candidate features for the machine-learning models. Summary ranges for *N*, FCR, NCPR, *κ*, and SCD for each subset are reported in Supplementary Table S1.

### 2.2 CALVADOS-2 simulations

All simulations were performed using the CALVADOS-2 coarse-grained model Tesei and Lindorff-Larsen (2023), in which each amino acid residue is represented by a single interaction site. Consecutive residues were connected by harmonic bonds. Nonbonded interactions were described by residue-specific Ashbaugh–Hatch short-range interactions and Debye–Hückel screened electrostatics, implemented as OpenMM CustomNonbondedForce terms. Salt concentration entered through the Debye screening length. Simulations were carried out using constant volume Langevin dynamics with the CUDA platform; full simulation parameters are reported in Table 1. Each sequence was simulated in the single chain dilute limit in a cubic periodic box of side length *L*_box_ = max(50, 4*N* ^0.6^) nm. The 50 nm lower bound was chosen to minimize interactions between periodic images, including for highly expanded conformations. Each of the 511 sequences was simulated independently at five monovalent salt concentrations (50, 100, 150, 300, and 500 mM), yielding a total of 2,555 single chain simulations. The 150 mM condition was simulated de novo for every sequence rather than reusing precomputed CALVADOS-2 trajectories from IDRome, ensuring that all five salt conditions were generated using a single internally consistent simulation protocol. Radius of gyration was monitored throughout each trajectory, and equilibrium ensemble averages were obtained from the production portion of the simulations. Convergence was assessed by block averaging of the *R*_g_ time series prior to downstream analysis.

**Table 1:** CALVADOS-2 simulation parameters. All 511 sequences were simulated at each of the five salt concentrations under an identical protocol.

| Parameter | Value |
| --- | --- |
| Force field | CALVADOS-2 Tesei and Lindorff-Larsen (2023) |
| Engine | OpenMM (CUDA platform) |
| Representation | One bead per residue |
| Electrostatics | Debye–Hückel (implicit monovalent salt) |
| Ensemble | NVT (Langevin thermostat) |
| Temperature | 293.15 K |
| pH | 7.5 |
| Salt concentrations | 50, 100, 150, 300, 500 mM |
| Box length | $\max(50, 4N^{0.6})$ nm, $N$ = sequence length |
| Saved frames | 1010 (first 10 discarded as equilibration) |
| Save interval | 7000 steps |
| Total steps | $7.07 \times 10^6$ per run |
| Total simulations | 2,555 (511 sequences × 5 salts) |

### 2.3 Observable extraction

For each trajectory, we extracted the radius of gyration *R*_g_ and end-to-end distance *R*_ee_ using the CALVADOS analysis workflow. The primary observable in this work is the ensemble-averaged radius of gyration,

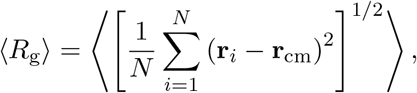

where **r**_*i*_ is the position of residue *i* and **r**_cm_ is the chain center of mass. The end-to-end distance was computed as

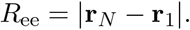

To quantify salt sensitivity, ensemble-averaged values of *R*_g_ were computed at each of the five simulated monovalent salt concentrations. The dependence of *R*_g_ on salt concentration was characterized by a linear fit to *R*_g_(*c*_salt_), and the resulting slope was used as the primary target variable in subsequent machine-learning analyses.

We also computed polymer shape descriptors from the extracted observables. In particular, we used dimensionless shape ratio

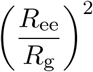

as a robust measure of the global chain shape.

An apparent scaling exponent *ν* was estimated from the internal-distance relation

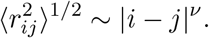

Because finite-chain effects and strongly collapsed or highly expanded conformations can lead to unstable estimates, *ν* was used only as a secondary structural descriptor and was not employed as a primary machine-learning target.

### 2.4 Salt-response fitting and regime classification

For each sequence, *R*_g_, *R*_ee_, and the ratio (*R*_ee_*/R*_g_)^2^ were fit as functions of monovalent salt concentration *c*_*s*_ (in molar units) across the five simulated salt points; sequences with fewer than four valid points for a given observable were excluded from that fit. The radius of gyration *R*_g_ was the primary observable for all subsequent analysis, while *R*_ee_ and (*R*_ee_*/R*_g_)^2^ were fit identically and retained as secondary diagnostics. The primary response coefficient was the slope of the linear fit

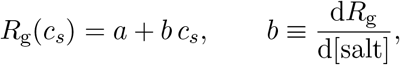

evaluated by least squares regression over all available salt points; positive *b* corresponds to salt-induced swelling and negative *b* to salt-induced contraction. A quadratic fit 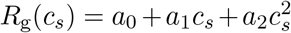 was also recorded as a diagnostic of curvature, but was not used for regime assignment. Each sequence was then assigned to one of four polymer physics regimes from the relative change in *R*_g_ across the sampled salt range and the shape of its discrete *R*_g_(*c*_*s*_) curve. We classified on the relative change rather than the fitted slope *b* in order to capture the total response magnitude without assuming linearity; it was defined from the simulated endpoints as

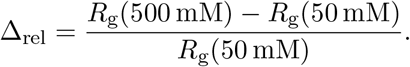

Sequences with |Δ_rel_ |*<* 0.03 were classified as *salt-insensitive*. Among the remaining sequences, a curve was deemed monotonic when its successive point-to-point differences in *R*_g_ were of a single sign (within a tolerance of 10^*−*3^ nm); monotonically decreasing curves were classified as *polyelectrolyte contraction* and monotonically increasing curves as *polyampholyte swelling*. Sequences whose *R*_g_(*c*_*s*_) curve was non-monotonic over the sampled range were classified as *non-monotonic*. Because this criterion operates directly on the discrete simulated values, the non-monotonic class is expected to be the most sensitive of the four to point wise statistical uncertainty in *R*_g_.

### 2.5 Polymer physics features

We used a compact set of eight sequence-derived descriptors motivated by charged-polymer theory and prior work on IDP sequence patterning. The features were selected to encode net charge, total charge content, charge patterning, chain length, and hydropathy while keeping the model low-dimensional and physically interpretable.

The net charge per residue was defined as

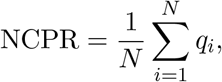

where *q*_*i*_ = +1 for Lys and Arg, *q*_*i*_ = − 1 for Asp and Glu, and *q*_*i*_ = 0 otherwise. The fraction of charged residues was defined as

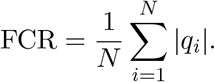

The sequence charge decoration was computed as

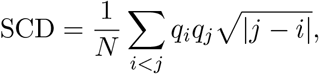

which captures the sequence-separation-weighted arrangement of charged residues along the chain. We also included the length-weighted quantity SCD × *N* to account for the extensive scaling of electrostatic patterning with chain length. Charge segregation was quantified using the Das–Pappu *κ* parameter. In addition, we included NCPR^2^ × *N*, FCR × *N*, *κ*FCR^2^, 1*/N*, and mean hydropathy as candidate predictors. The final feature set is summarized in Table 2.

**Table 2:** Polymer-physics descriptors used as machine-learning features.

| Feature | Description | Motivation |
| --- | --- | --- |
| ncpr_sq_length | $\text{NCPR}^2 \times N$ | net-charge scaling |
| scd | sequence charge decoration | charge patterning |
| scd_length | $\text{SCD} \times N$ | length-weighted patterning |
| fcr_length | $\text{FCR} \times N$ | total charge content |
| kappa_fcr_sq | $\kappa \times \text{FCR}^2$ | charge segregation at finite charge fraction |
| inv_length | $1/N$ | finite-size correction |
| mean_hydropathy | Kyte–Doolittle average | residue-specific solvation |
| kappa | charge-patterning parameter | charge segregation |

### 2.6 Machine learning

We trained models to predict the salt-response slope *b* = d*R*_g_*/*d[salt] from the eight polymer-physics descriptors. As an interpretable linear baseline we used ridge regression on standardized features (*α* = 1.0). To capture nonlinear descriptor response relationships we trained a gradient-boosted regressor (XGBoost) with 500 trees, maximum depth 4, learning rate 0.05, and subsample and column subsample fractions of 0.85; tree models were trained on raw (unstandardized) features, to which they are invariant. The patterning parameter *κ*, undefined for the pure-polyelectrolyte sequences of a single charge sign, was set to zero for those sequences; *κ* is among the least important features (Section 3.4).

Regression performance was assessed with two strategies on identical folds. Random five fold cross validation, splitting sequences into folds at random, estimates in distribution interpolation. Leave-one-subset-out cross-validation, holding out each of the four library subsets in turn, assesses generalization across design classes. Performance was quantified by the coefficient of determination *R*^2^ and the mean absolute error (MAE); we report both because *R*^2^ is uninformative for the salt-insensitive regime, whose near-constant slopes give negligible target variance.

For regime classification we predicted the four salt response classes from the same eight descriptors using a logistic-regression baseline (on standardized features) and a gradient-boosted (XGBoost) classifier (200 trees, maximum depth 4, learning rate 0.10, subsample and column-subsample 0.85, multinomial softmax objective). Because the regimes are imbalanced, performance was evaluated by stratified five fold cross-validation preserving class proportions, and summarized by overall accuracy, per-class *F*_1_, and the confusion matrix.

To interpret the nonlinear models, we computed SHAP values using the TreeExplainer algorithm. We report global feature importance as the mean absolute SHAP value per feature, and SHAP dependence plots for the dominant descriptors to expose nonlinear feature response relationships.

## 3 Results

### 3.1 Salt response varies dramatically across the library

Across the 511 sequences in the library, the radius of gyration responds to monovalent salt in qualitatively different ways. The slope d*R*_g_*/*d[salt] ranges from −4.12 to +5.16 nm/M, wide enough that added salt compacts some chains and expands others by comparable amounts. We grouped sequences into four regimes from the shape of the *R*_g_–[salt] curve and its relative change between 50 and 500 mM (Fig. 2): polyelectrolyte-like contraction (PE contraction, *n* = 82), polyampholyte-like swelling (PA swelling, *n* = 187), non-monotonic (*n* = 117), and salt-insensitive (*n* = 125). The two directional regimes set the extremes. The most strongly contracting sequences, taken from the NCPR series, lose about 28% of their 50 mM size by 500 mM, and the strongest expanders, well-mixed *κ*-variants, gain about 78% over the same range. Non-monotonic chains reverse direction within the window, and salt-insensitive chains change little at any concentration.

**Figure 1:**
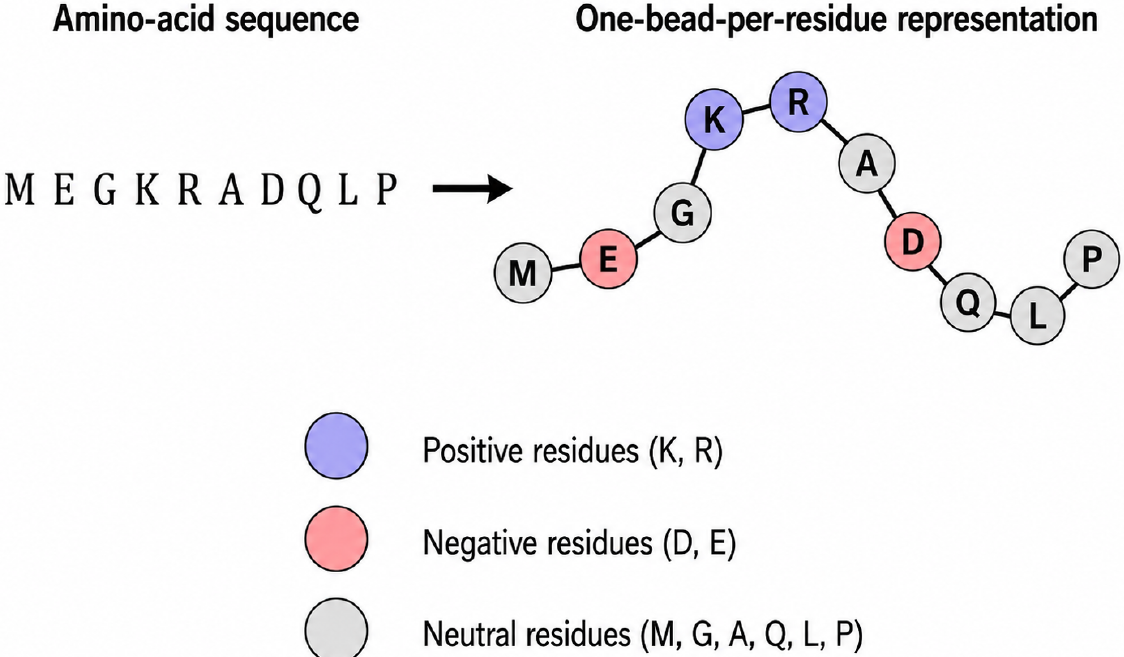
Schematic representation of the CALVADOS-2 coarse-grained model. An amino-acid sequence is mapped to a one-bead-per-residue polymer representation, in which each residue is represented by a single interaction site connected along the chain backbone. Beads are colored here by residue charge class for illustration: positively charged residues (K, R), negatively charged residues (D, E), and neutral residues.

**Figure 2:**
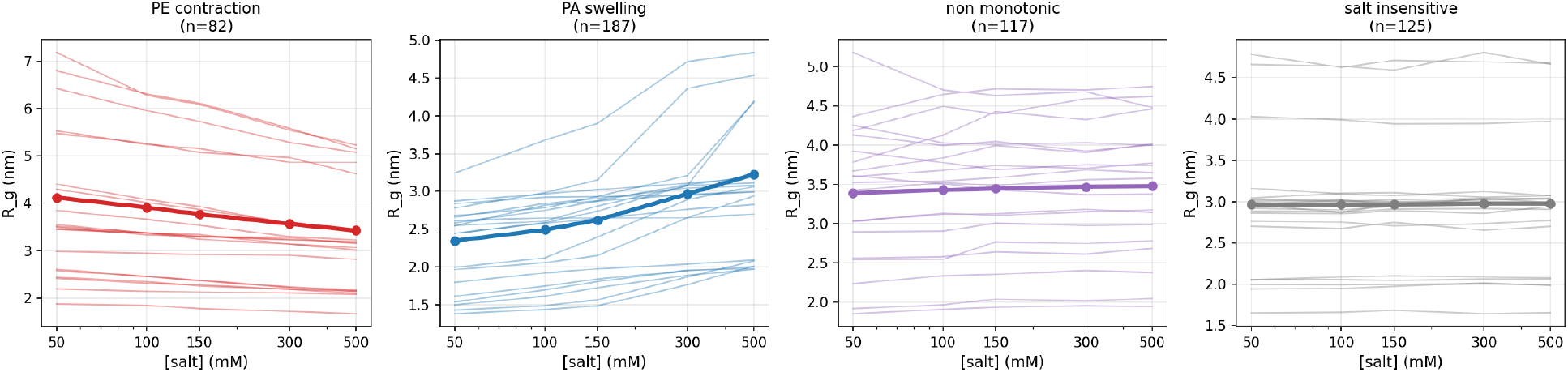
Salt-response curves by polymer-physics regime. Each panel shows ~20 representative sequences (thin lines) and the ensemble mean (thick line with markers) for one regime, with sample sizes in panel titles. Regimes are classified by the shape of the *R*_g_-vs-[salt] curve and the relative change from 50 to 500 mM: PE contraction and PA swelling are the monotonic directional classes, non-monotonic chains reverse direction within the window, and salt-insensitive chains change little across the range.

### 3.2 Regimes cluster in sequence space along SCD

The four regimes separate along a single sequence coordinate, the sequence charge decoration SCD (Fig. 3). PA-swelling sequences fall in the negative tail (SCD < −0.5), PE-contraction sequences in the positive tail (SCD *>* +1), and non-monotonic sequences in the low-magnitude middle (|SCD| *<* 1). The directional regimes barely overlap in SCD even though both span the full range of chain lengths in the library, which says that charge patterning rather than size is the dominant axis associated with the sign of the response. Salt-insensitive sequences are the exception: rather than occupying a band of their own they spread across the whole SCD axis. Their shared feature is a low fraction of charged residues, so there is little electrostatic interaction for salt to screen however the few charges are arranged.

**Figure 3:**
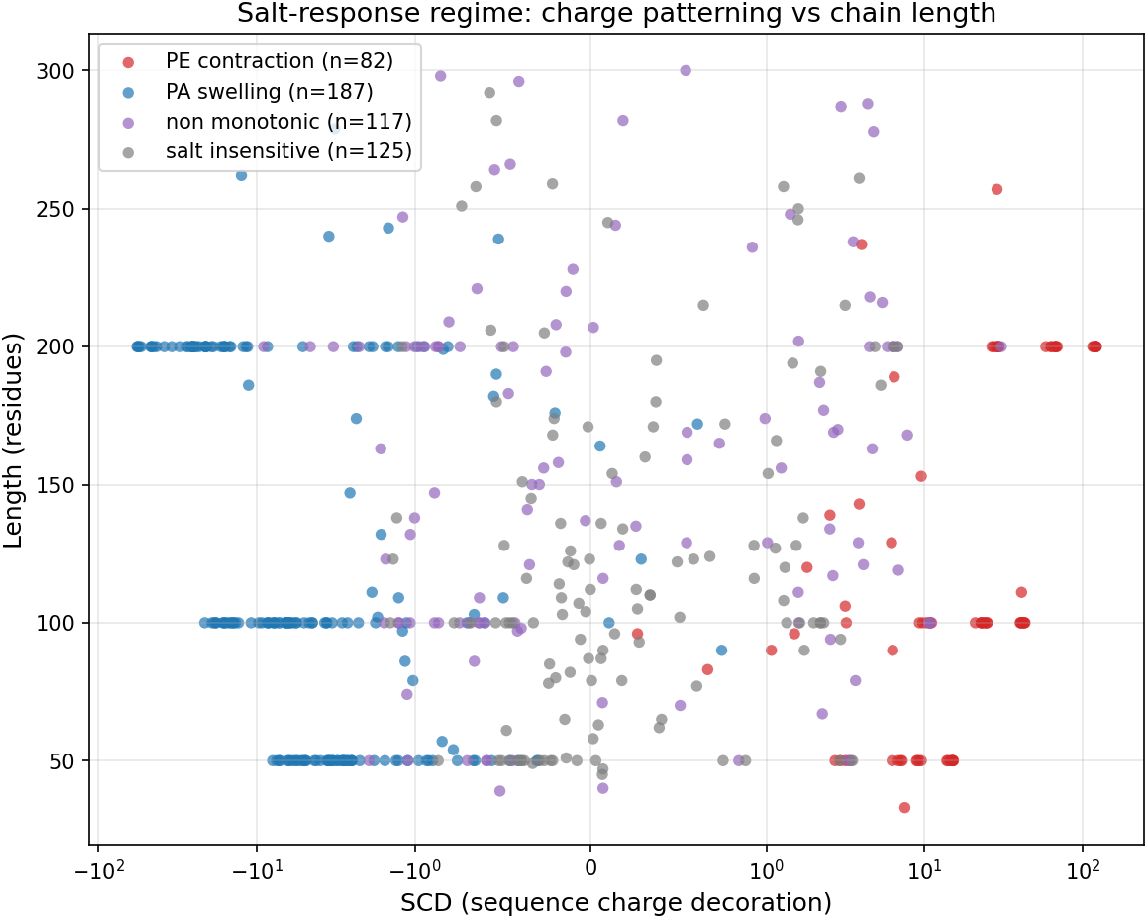
Polymer-physics regimes in sequence-charge-decoration (SCD) versus chain-length space. Each point is one of 511 sequences, coloured by salt-response regime (SCD on a symlog axis). The directional regimes separate along SCD with little overlap and across all chain lengths, identifying charge patterning rather than size as the dominant axis of salt-response variation; salt-insensitive sequences spread across SCD and are instead set apart by a low fraction of charged residues.

The split has a direct electrostatic reading. Sequences with well-mixed opposite charges (the polyampholyte end) are held compact by intra-chain attraction at low salt; screening that attraction lets the chain swell. Sequences dominated by like-charge repulsion, often those with substantial net charge, are expanded by intrachain repulsion at low salt; screening that repulsion lets the chain contract. The sign of SCD tracks which of the two dominates.

### 3.3 Polymer-physics features predict salt-response slope

Eight polymer-physics descriptors predict the salt-response slope with useful accuracy. Under random 5-fold cross-validation, ridge regression reaches *R*^2^ = 0.832 ± 0.029 (MAE 0.44 nm/M) and gradient-boosted trees (XGBoost) reach *R*^2^ = 0.966 ± 0.011 (MAE 0.18 nm/M) on the same features (Fig. 4). The ridge residuals are largest at the contracting and expanding tails, where a linear fit pulls the extremes toward the mean, whereas XGBoost tracks both tails.

**Figure 4:**
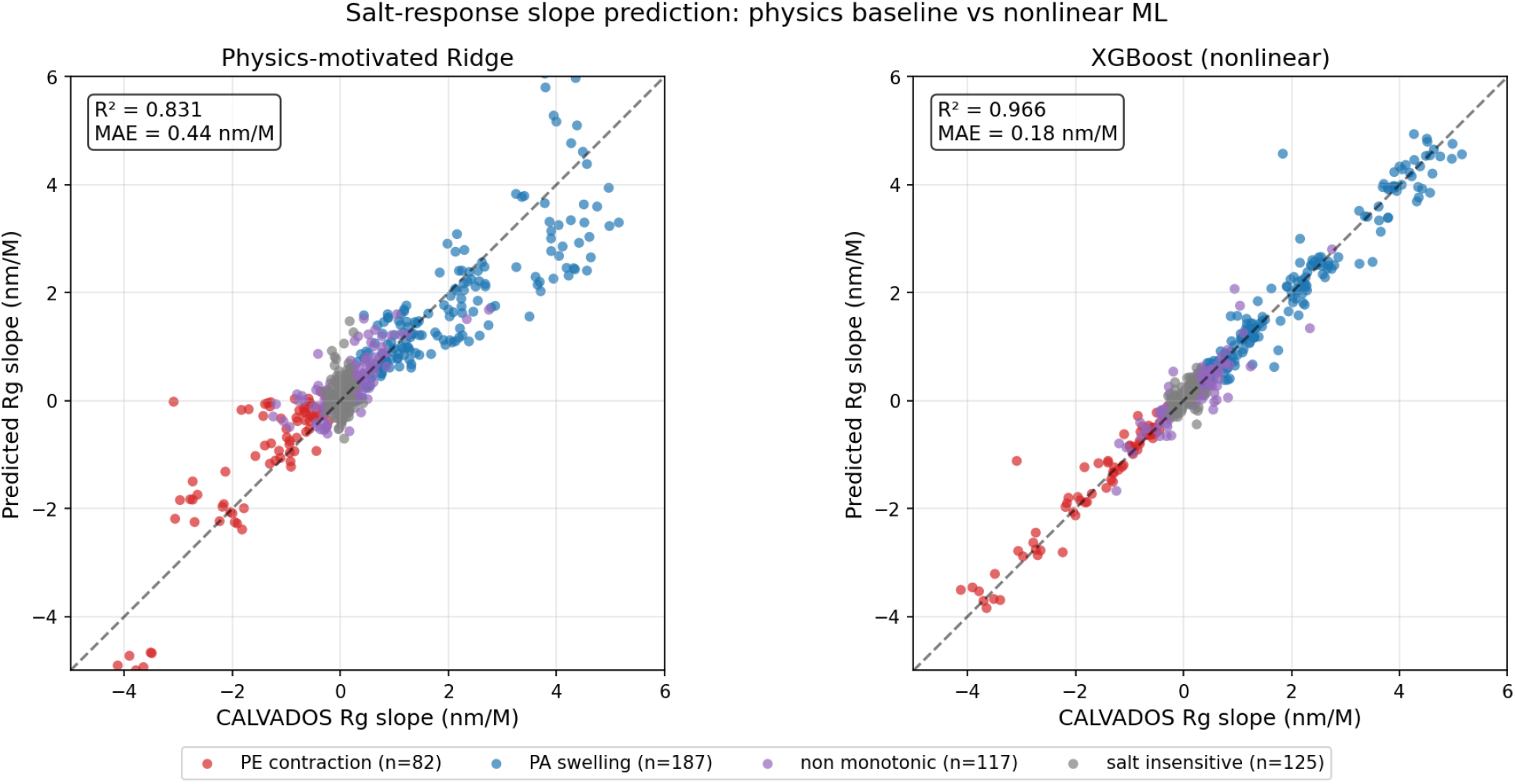
Predicted versus simulated salt-response slope (d*R*_g_*/*d[salt], nm/M) under random 5-fold cross-validation. Left: physics-motivated ridge regression on 8 descriptors (*R*^2^ = 0.83, MAE 0.44 nm/M). Right: XGBoost on the same features (*R*^2^ = 0.97, MAE 0.18 nm/M). Points are coloured by polymer-physics regime; the dashed line is *y* = *x*. Under leave-one-subset-out cross-validation the same models reach mean *R*^2^ = −0.13 (ridge) and 0.60 (XGBoost); see text.

A second, harder number matters for how these models should be used. The library is built from distinct sequence families (NCPR series, *κ* variants, and others), so random cross-validation mixes near-relatives across folds and measures interpolation within a family. Leave-one-subset-out (LOSO) cross-validation instead holds out whole families and measures extrapolation to sequence types the model has not seen. Under LOSO the ridge model loses predictive power (mean *R*^2^ = − 0.13, worse than predicting the library mean), whereas XGBoost retains moderate transferability (*R*^2^ = 0.60).

The two scores answer different questions: random CV is the in-distribution accuracy, LOSO is the deployment-relevant estimate for genuinely new sequences. We report both, and read the gap as the caution that applies before using either model outside the families sampled here.

### 3.4 SHAP analysis identifies SCD-length as the master predictor

To see which descriptors carry the prediction and how, we computed SHAP attributions for the XGBoost regressor (Fig. 5). One feature dominates: the product of SCD and chain length, SCD × *N*, with mean |SHAP| of 0.643 nm/M, more than twice the next feature (SCD alone, 0.290) and well above NCPR^2^ ×*N* (0.152), FCR × *N* (0.120), *κ* FCR^2^ (0.101), *κ* (0.094), 1*/N* (0.056), and mean hydropathy (0.029). The attribution depends on its own value as a sigmoid, not a line (Fig. 5a): it saturates near +2.2 nm/M for strongly polyampholytic sequences (SCD × *N* ≪ 0), crosses zero near SCD × *N* ≈ 0, and saturates near −1.8 nm/M for strongly polyelectrolytic sequences (SCD × *N* ≫ 0). The sign of SCD × *N* therefore sets the direction of the salt response and its magnitude sets the strength, the same threshold behaviour seen in the regime map of Section 3.2. This sigmoidal dependence explains why XGBoost outperforms ridge: a linear model cannot saturate, so it overshoots in the tails and undershoots near the threshold.

**Figure 5:**
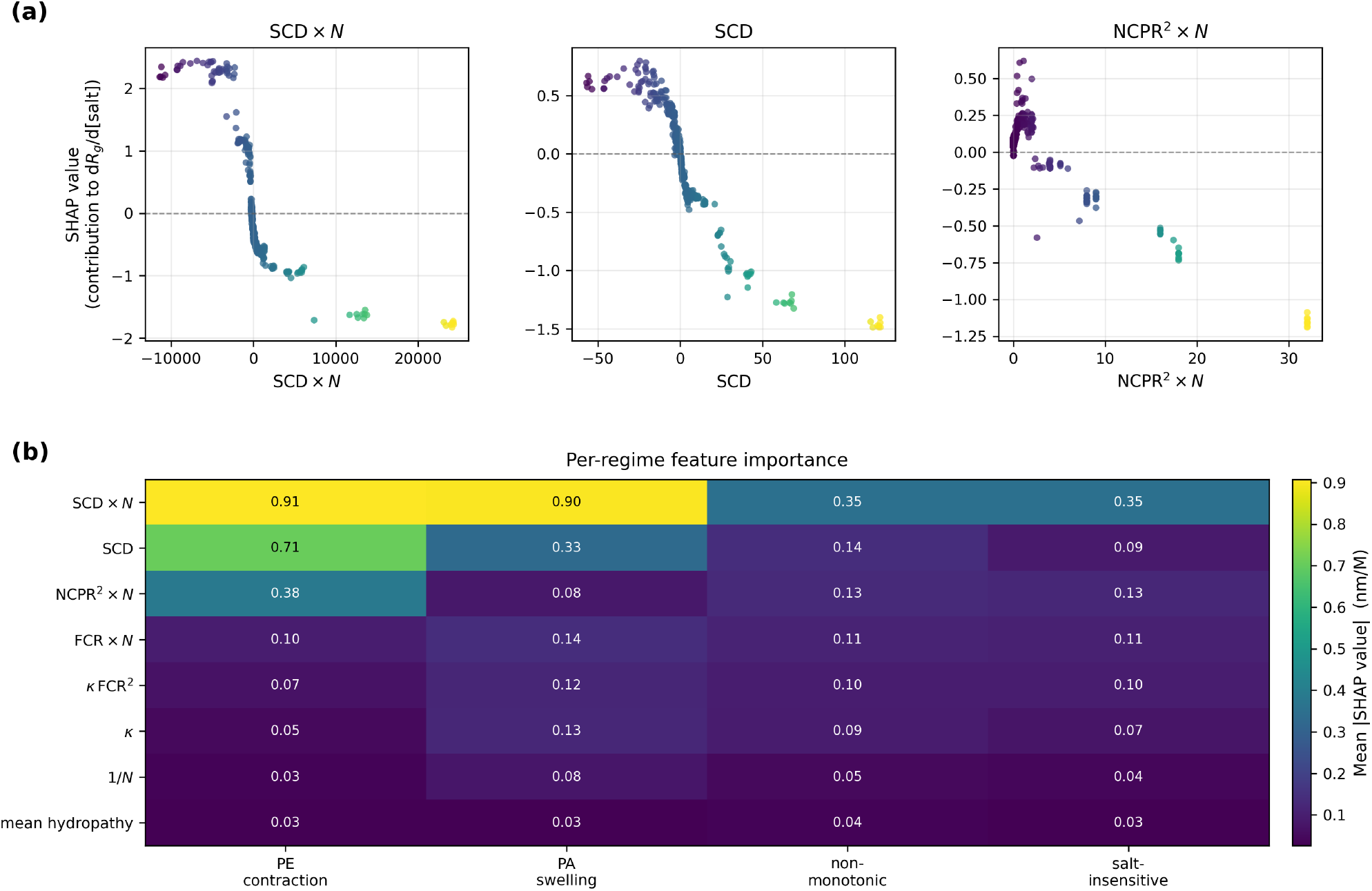
Interpretable attribution of the predicted salt-response slope d*R*_g_*/*d[salt] via SHAP analysis of the gradient-boosted model. (a) SHAP dependence plots for the three highest-attribution features. The contribution of SCD × *N* (left) is sigmoidal: it saturates near +2.2 for strongly polyampholytic sequences (SCD × *N* ≪ 0), passes through zero near SCD × *N* ≈ 0, and saturates near −1.8 for strongly polyelectrolytic sequences (SCD × *N* ≫ 0), so that the sign of SCD × *N* sets the direction of salt response and its magnitude sets the strength. SCD (centre) and NCPR^2^ × *N* (right) contribute in the same sense but with smaller magnitude. (b) Mean |SHAP| per feature within each regime. SCD × *N* is the dominant coordinate in both directional regimes (0.91 in PE contraction, 0.90 in PA swelling), with additional contributions from SCD (0.71) and NCPR^2^ × *N* (0.38) in PE contraction; in the non-monotonic and salt-insensitive regimes no single feature dominates (all mean |SHAP| ≤ 0.35), consistent with these regimes being poorly separated by static sequence descriptors. Colour in (a) encodes feature value; colour in (b) encodes mean |SHAP| (nm/M).

Within regimes (Fig. 5b), SCD × *N* leads both directional classes (mean |SHAP| 0.91 for PE contraction, 0.90 for PA swelling), with SCD (0.71) and NCPR^2^ × *N* (0.38) adding signal in the contraction class. In the non-monotonic and salt-insensitive classes no feature exceeds 0.35, consistent with these regimes being poorly separated by any static sequence descriptor. Mean hydropathy is the weakest contributor by an order of magnitude, which is expected for CALVADOS-2: its salt dependence enters only through a Debye-Hückel screening term on the charged residues, with no salt coupling to hydrophobic contacts. Thus, the feature attribution reflects the salt-dependent physics encoded in the force field.

### 3.5 Regime classification: direction is predictable, magnitude harder

As a four way classification problem the picture is the same but sharper: the direction of the salt response is easy to predict, its finer structure is not. The XGBoost classifier reaches 73% accuracy under 5-fold cross validation (Fig. 6), well above the 37% majority class baseline, and its balanced accuracy is 0.71, so the class imbalance does not inflate the headline figure. Performance falls into two tiers. The directional regimes are recovered cleanly (PA swelling *F*_1_ = 0.87, recall 0.88; PE contraction *F*_1_ = 0.81, recall 0.80), while the others trail (salt-insensitive 0.69, non-monotonic 0.46).

**Figure 6:**
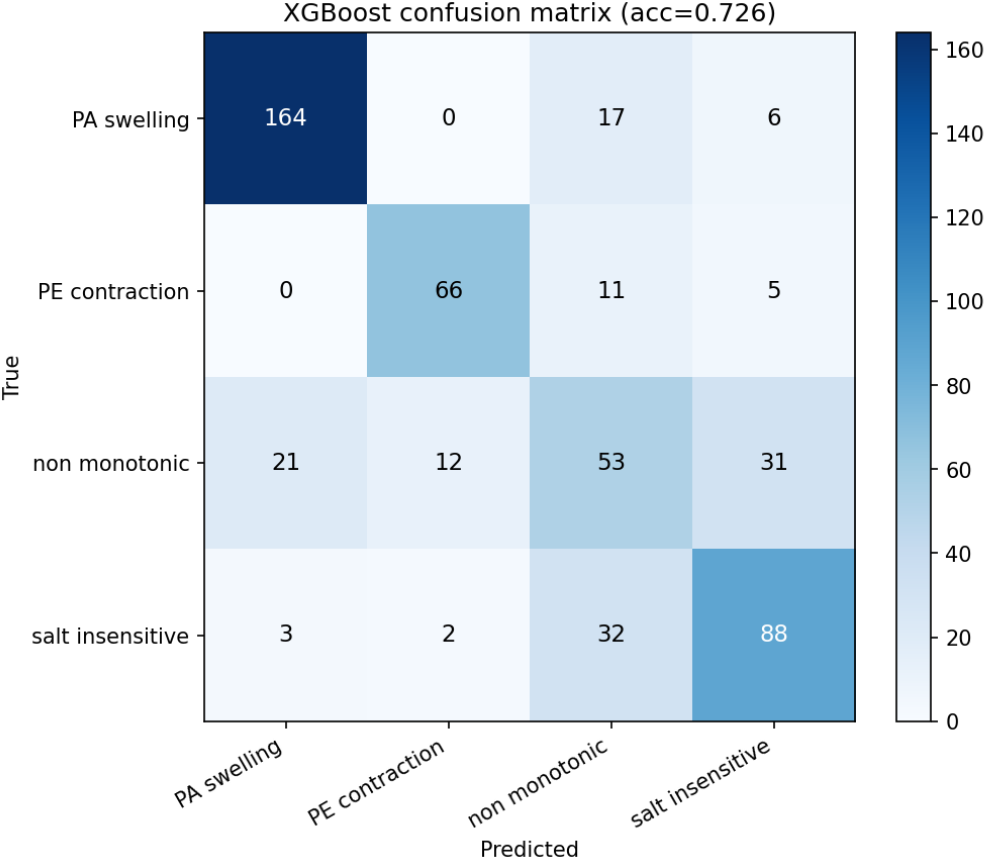
Confusion matrix for the XGBoost regime classifier (5-fold cross-validation, overall accuracy 0.73). The zero off-diagonal between PA swelling and PE contraction shows that the direction of salt response is robustly predictable; errors concentrate between non-monotonic and salt-insensitive, both of which have small slope magnitudes and are distinguished only by curve shape at the five salt points sampled

The clearest signal is an empty cell. No swelling chain in the cross-validation was labelled contracting, and none in the reverse direction, so the two directional classes are never confused with each other: the sign of the response is encoded robustly by the charge features. The non-monotonic class is the weakest and is confused in every direction, losing 31 of its 117 members to salt-insensitive, 21 to PA swelling, and 12 to PE contraction (recall 0.45, precision 0.47). Its largest single overlap is with salt-insensitive and is roughly symmetric (31 one way, 32 the other). Both classes carry small slope magnitudes and differ mainly in curve shape, which five salt points (50–500 mM) resolve only coarsely; separating a flat chain from a weakly non-monotonic one would likely benefit from finer salt sampling rather than additional sequence features.

## 4 Discussion

SHAP attribution identifies SCD × *N* as the dominant coordinate governing salt response across the library, accounting for roughly 40% of total feature attribution and exceeding the next feature more than twice. This assigns quantitative, library-scale weight to a descriptor that polyampholyte theory had identified for IDR dimensions Lin et al. (2016); Sawle and Ghosh (2015), extending its role to the salt-response slope specifically. The attribution is sigmoidal in SCD × *N*, saturating for strongly polyampholytic (SCD × *N* ≪ 0) and polyelectrolytic (SCD × *N* ≫ 0) sequences with a sharp crossover near zero, so that the sign of SCD × *N* sets the direction of response and its magnitude the strength. This crossover behaviour matches the random-phase-approximation prediction that polyelectrolytic and polyampholytic sequences respond to salt in opposite directions Huihui et al. (2018), and is precisely the structure a linear model cannot represent, explaining why the gradient-boosted model captures it while the ridge baseline does not.

The two directional regimes are well described by these descriptors: both are predicted with per-regime *R*^2^ *>* 0.9 under random cross-validation, and the classifier separates them with per class *F*_1_ of 0.81 and 0.87 and no PE–PA confusion. The non-monotonic and salt-insensitive regimes are harder, for reasons that are informative rather than technical: both are defined by properties static charge descriptors do not encode directly, curve shape and the near-absence of response, respectively, and SHAP confirms that no single feature dominates either (|SHAP| ≤ 0.35 throughout), in contrast to the *>* 0.90 of SCD × *N* in the directional regimes. The salt-insensitive class is moreover heterogeneous, spanning near-neutral, charge-balanced, and weakly charged hydrophobic sequences, so a single coordinate is not expected to organize it. Classifier errors accordingly concentrate between these two small slope regimes.

These results complement sequence-conditioned ensemble models. Predictors such as ALBA-TROSS Lotthammer et al. (2024) and generative models such as STARLING Novak et al. (2026) supply ensemble properties at a specified ionic strength; our contribution is orthogonal, identifying which interpretable descriptors control the *response* to changing salt and organizing sequences into regimes. Where theory predicts salt response for individual sequences Lin et al. (2016); Huihui et al. (2018), we show the same physical quantities emerge empirically as the dominant predictors across hundreds of sequences; where coarse-grained surveys characterized the mechanistic ingredients of salt response Wohl et al. (2021), we convert that understanding into a predictive sequence-to-regime map.

Several limitations bound these conclusions. CALVADOS-2 represents salt only through Debye– Hückel screening and omits residue-specific salting-out and Hofmeister effects Wohl et al. (2021); responses driven by such mechanisms are outside its scope, a caveat most relevant to the weakly charged low-FCR subset. The feature set was also restricted to composition- and patterning-derived descriptors computable from sequence alone; the salt-specific extensions SCD_lowsalt_ and SCD_highsalt_ introduced by Huihui et al. (2018) were not included as candidate predictors. Whether those descriptors, which were designed specifically for salt-response prediction, would displace or complement SCD × *N* in the SHAP ranking is a direct open question. The random-CV performance (*R*^2^ = 0.97) is optimistic, since compositionally related sequences leak across folds; the leave-one-subset-out value (*R*^2^ = 0.60) is the honest estimate for unseen sequence families and the deployment-relevant metric. For the salt-insensitive regime, *R*^2^ is uninformative given the near-constant slopes, and the mean absolute error (0.11 nm/M, the lowest of any regime) is the appropriate measure.

Finally, the study is entirely simulation-based, and experimental benchmarking of the predicted slopes is the natural next step. The salt-response slope d*R*_g_*/*d[salt] is accessible to established solution-scattering and single-molecule techniques: small-angle X-ray scattering, single-molecule Förster resonance energy transfer, and size-exclusion chromatography all resolve IDP dimensions as a function of ionic strength, and salt-dependent measurements of this kind underpin several of the studies motivating this work Müller-Späth et al. (2010); Waszkiewicz et al. (2024).We therefore frame the predicted slopes as directional and relative rather than quantitatively calibrated, since CALVADOS-2 is parameterized to reproduce ensemble dimensions near physiological ionic strength and its absolute salt dependence has not been independently benchmarked. A further constraint is the salt sampling itself: five concentrations spanning 50 mM to 500 mM resolve the *shape* of *R*_g_(*c*_*s*_) only coarsely, which is the principal reason the non-monotonic and salt-insensitive classes remain difficult to separate.

## Acknowledgements

Computational resources were provided by the Center for Computationally Assisted Science and Technology (CCAST) at North Dakota State University. ChatGPT was used as an assistant tool for text polishing and proofreading. The author has reviewed the content of this publication and takes full responsibility for its findings.

## Data availability

### Underlying data

Sequence libraries, simulation input files, processed observables, salt-response fits, and analysis scripts required to reproduce the results presented in this work are available at https://github.com/Aryal-Mahesh/idp-salt-response.

## Notes

### Competing Interest Statement

The authors have declared no competing interest.

### Summary of Updates

Revised version. The Introduction has been restructured to frame IDP salt response within the broader polymer-physics context of chain conformational response to solvent environment, with additional references to foundational excluded-volume, depletion, and polyelectrolyte theory. The Discussion has been expanded: the limitations section now addresses the omission of salt-specific SCD variants as candidate predictors, the accessibility of the predicted salt-response slopes to solution-scattering and single-molecule measurement, and the constraint imposed by the five-point salt sampling on resolving curve shape. No changes were made to the simulation data, machine-learning results, figures, or numerical findings.

## References

Sho Asakura and Fumio Oosawa. On interaction between two bodies immersed in a solution of macromolecules. Journal of Chemical Physics, 22(7):1255–1256, 1954. doi: 10.1063/1.1740347.

Pierre-Gilles de Gennes. Scaling Concepts in Polymer Physics. Cornell University Press, Ithaca, NY, 1979.

Michael Rubinstein and Ralph H. Colby. Polymer Physics. Oxford University Press, New York, 2003.

Andrey V. Dobrynin, Ralph H. Colby, and Michael Rubinstein. Scaling theory of polyelectrolyte solutions. Macromolecules, 28(6):1859–1871, 1995. doi: 10.1021/ma00110a021.

Mahesh Aryal and Alan R. Denton. Response of hydrogels and microgels to nanoparticle crowding. Journal of Chemical Physics, 164(6):064904, 2026. doi: 10.1063/5.0312584.

Peter E. Wright and H. Jane Dyson. Intrinsically disordered proteins in cellular signalling and regulation. Nature Reviews Molecular Cell Biology, 16(1):18–29, 2015. doi: 10.1038/nrm3920.

Robin van der Lee, Marija Buljan, Benjamin Lang, Robert J. Weatheritt, Gary W. Daughdrill, A. Keith Dunker, Monika Fuxreiter, Julian Gough, Joerg Gsponer, David T. Jones, Philip M. Kim, Richard W. Kriwacki, Christopher J. Oldfield, Rohit V. Pappu, Peter Tompa, Vladimir N. Uversky, Peter E. Wright, and M. Madan Babu. Classification of intrinsically disordered regions and proteins. Chemical Reviews, 114(13):6589–6631, 2014. doi: 10.1021/cr400525m.

Sonja Müller-Späth, Andrea Soranno, Verena Hirschfeld, Hagen Hofmann, Stefan Rüegger, Luc Reymond, Daniel Nettels, and Benjamin Schuler. Charge interactions can dominate the dimensions of intrinsically disordered proteins. Proceedings of the National Academy of Sciences, 107(33): 14609–14614, 2010.

Radost Waszkiewicz, Agnieszka Michaś Michał K. Białobrzewski, Barbara P. Klepka, Maja K. Cieplak-Rotowska, Zuzanna Staszałek, Bogdan Cichocki, Maciej Lisicki, Piotr Szymczak, and Anna Niedzwiecka. Hydrodynamic radii of intrinsically disordered proteins: Fast prediction by minimum dissipation approximation and experimental validation. Journal of Physical Chemistry Letters, 15(19):5024–5033, 2024. doi: 10.1021/acs.jpclett.4c00312.

Jasmine Cubuk and Andrea Soranno. Macromolecular crowding and intrinsically disordered proteins: A polymer physics perspective. ChemSystemsChem, 4(4):e202100051, 2022. doi: 10.1002/syst.202100051.

Austin Banks, Sanbo Qin, Kevin L. Weiss, Christopher B. Stanley, and Huan-Xiang Zhou. Intrinsically disordered protein exhibits both compaction and expansion under macromolecular crowding. Biophysical Journal, 114(5):1067–1079, 2018. doi: 10.1016/j.bpj.2018.01.011.

Yi-Hsuan Lin, Julie D. Forman-Kay, and Hue Sun Chan. Sequence-specific polyampholyte phase separation in membraneless organelles. Physical Review Letters, 117(17):178101, 2016. doi: 10.1103/PhysRevLett.117.178101.

Basile I. M. Wicky, Sarah L. Shammas, and Jane Clarke. Affinity of idps to their targets is modulated by ion-specific changes in kinetics and residual structure. Proceedings of the National Academy of Sciences of the United States of America, 114(37):9882–9887, 2017. doi: 10.1073/pnas.1705105114.

Tanja Mittag and Rohit V. Pappu. A conceptual framework for understanding phase separation and addressing open questions and challenges. Molecular Cell, 82(12):2201–2214, 2022. doi: 10.1016/j.molcel.2022.05.018.

Paul G. Higgs and Jean-François Joanny. Theory of polyampholyte solutions. Journal of Chemical Physics, 94(2):1543–1554, 1991. doi: 10.1063/1.460012.

Rahul K. Das and Rohit V. Pappu. Conformations of intrinsically disordered proteins are influenced by linear sequence distributions of oppositely charged residues. Proceedings of the National Academy of Sciences of the United States of America, 110(33):13392–13397, 2013. doi: 10.1073/pnas.1304749110.

Lucas Sawle and Kingshuk Ghosh. A theoretical method to compute sequence dependent configurational properties in charged polymers and proteins. Journal of Chemical Physics, 143(8):085101, 2015. doi: 10.1063/1.4929391.

Jonathan Huihui, Taylor Firman, and Kingshuk Ghosh. Modulating charge patterning and ionic strength as a strategy to induce conformational changes in intrinsically disordered proteins. Journal of Chemical Physics, 149(8):085101, 2018. doi: 10.1063/1.5037727.

Samuel Wohl, Matthew Jakubowski, and Wenwei Zheng. Salt-dependent conformational changes of intrinsically disordered proteins. Journal of Physical Chemistry Letters, 12:6684–6691, 2021. doi: 10.1021/acs.jpclett.1c01607.

Giulio Tesei, Anna Ida Trolle, Nicolas Jonsson, Johannes Betz, Frederik E. Knudsen, Francesco Pesce, Kristoffer E. Johansson, and Kresten Lindorff-Larsen. Conformational ensembles of the human intrinsically disordered proteome. Nature, 626(8000):897–904, 2024. doi: 10.1038/s41586-023-07004-5.

Jeffrey M. Lotthammer, Garrett M. Ginell, Daniel Griffith, Ryan J. Emenecker, and Alex S. Holehouse. Direct prediction of intrinsically disordered protein conformational properties from sequence. Nature Methods, 21:465–476, 2024. doi: 10.1038/s41592-023-02159-5.

Borna Novak, Jeffrey M. Lotthammer, Ryan J. Emenecker, and Alex S. Holehouse. Accurate predictions of disordered protein ensembles with starling. Nature, 652(8108):240–250, 2026. doi: 10.1038/s41586-026-10141-2.

Alex S. Holehouse, Rahul K. Das, James N. Ahad, Mary O. G. Richardson, and Rohit V. Pappu. Cider: resources to analyze sequence-ensemble relationships of intrinsically disordered proteins. Biophysical Journal, 112(1):16–21, 2017. doi: 10.1016/j.bpj.2016.11.3200.

Jack Kyte and Russell F. Doolittle. A simple method for displaying the hydropathic character of a protein. Journal of Molecular Biology, 157(1):105–132, 1982. doi: 10.1016/0022-2836(82)90515-0.

Giulio Tesei and Kresten Lindorff-Larsen. Improved predictions of phase behaviour of intrinsically disordered proteins by tuning the interaction range. Open Research Europe, 2:94, 2023. doi: 10.12688/openreseurope.14967.2.

