## Supplementary material for "Sequence charge decoration organizes salt response regimes in intrinsically disordered proteins: an interpretable machine-learning study": Composition and charge pattering ranges of sequence library subset (from another version of this preprint: version 1, 2026-06-04)

Mahesh Aryal

June 1, 2026

This Supplementary Information provides composition and charge-patterning ranges for the sequence subsets used in the main text.

### Supplementary Tables

Table S1: Composition and charge-patterning ranges of the four sequence-library subsets. Values are reported as min–max across sequences in each subset; the NCPR-series subset was designed at fixed FCR = 0.40. $N$, sequence length; FCR, fraction of charged residues; NCPR, net charge per residue; $\kappa$, charge-patterning parameter [1]; SCD, sequence charge decoration [2].

| Subset | Count | $N$ | FCR | NCPR | $\kappa$ | SCD |
|---|---|---|---|---|---|---|
| $\kappa$-variants | 195 | 50 to 200 | 0.30 to 0.50 | −0.02 to 0.00 | 0.04 to 0.59 | −56.89 to −0.22 |
| NCPR series | 108 | 50 to 200 | 0.40 | −0.40 to 0.40 | 0.07 to 0.41 | −1.43 to 121.62 |
| IDRome (stratified) | 150 | 33 to 300 | 0.04 to 0.59 | −0.42 to 0.19 | 0.09 to 0.42 | −12.45 to 41.02 |
| IDRome (low-FCR) | 58 | 47 to 298 | 0.04 to 0.10 | −0.04 to 0.07 | 0.13 to 0.36 | −0.85 to 4.75 |
| **Full library** | 511 | 33 to 300 | 0.04 to 0.59 | −0.42 to 0.40 | 0.04 to 0.59 | −56.89 to 121.62 |

### References

[1] Rahul K. Das and Rohit V. Pappu. Conformations of intrinsically disordered proteins are influenced by linear sequence distributions of oppositely charged residues. *Proceedings of the National Academy of Sciences of the United States of America*, 110(33):13392–13397, 2013. doi: 10.1073/pnas.1304749110.

[2] Lucas Sawle and Kingshuk Ghosh. A theoretical method to compute sequence dependent configurational properties in charged polymers and proteins. *Journal of Chemical Physics*, 143(8): 085101, 2015. doi: 10.1063/1.4929391.
